# Co-existence of ST-Segment Depression and Exaggerated Blood Pressure Response in Older Runners

**DOI:** 10.1101/2024.10.29.620598

**Authors:** Kazumasa Manabe, Owen N. Beck, Justus D. Ortega, Hirofumi Tanaka

**Author notes:** **Correspondence:** Hirofumi Tanaka., Ph.D., Department of Kinesiology and Health Education, University of Texas at Austin, 2109 San Jacinto Blvd D3700, Austin, TX 78712, USA. **Sources of support:** Kazumasa Manabe is currently receiving a research fellowship for young scientists (23KJ1618) from the Japan Society for the Promotion of Science.

## Abstract

Exercise stress tests are typically performed on symptomatic patients and those at elevated risks of ischemic heart disease. In more recent years, these tests are increasingly recommended for masters athletes who regularly perform exercise training. In the sample of apparently healthy physically active older adults who were recruited for a research study to evaluate exercise economy, we found the coexistence of ST-segment depression and exaggerated BP response in older runners. These findings are consistent with the notion that the screening and assessment of cardiovascular risks should be extended to masters athletes who vigorously train regularly and are seemingly healthy.

## INTRODUCTION

Treadmill exercise stress test on electrocardiogram (ECG) is one of the most frequently-used noninvasive modalities for screening ischemic heart disease. ST-segment depression during exercise is a significant marker that is linked to myocardial ischemia ^1,2^. Additionally, peak systolic blood pressure (BP) is another marker that is associated with adverse cardiac events ^3^. In a study using ambulatory ECG and BP monitoring, transient ischemia is often presented with both ST-segment depression and increases in BP ^4^. Exercise stress tests are typically conducted in symptomatic patients and those who are at higher risk of ischemic heart disease. In more recent years, the recommendation to screen and assess cardiovascular risks has extended to a variety of populations including chronically active and seemingly healthy older athletes ^1,2^.

Regular exercise is known to confer substantial benefits for cardiovascular health. Even in apparently healthy athletes, exercise stress tests are conducted to gauge the risk of strenuous training and competition and to monitor fitness and conditioning ^1,2^. The interpretation of ST segment depression and peak systolic BP can be more nuanced in endurance-trained athletes ^1,5,6^. In the retrospective analysis of a previous research study to evaluate walking economy in older adults ^7^, we observed a fairly high incidence of both ST segment depression and peak systolic BP in older endurance-trained athletes. Herein we report such observations.

## METHODS

We retrospectively analyzed prior data involving apparently healthy older adults (aged ≥65 years) who engaged in regular running (12 men and 5 women) and in walking for exercise (3 men and 18 women) ^7^. All the participants self-reported engaging in their respective activities three or more times per week for at least 30 minutes per session for at least six months prior to the study. None of the participants had been diagnosed with neurological, orthopedic, or cardiovascular disorders before the test. All participants completed ECG at rest and during a maximal treadmill test to determine maximal aerobic capacity, with BP measured using auscultation. A physician interpreted the ECG records. ST-segment depression was defined as ST-segment >1 mm lower than the baseline ^2^. All participants provided written informed consent, and the study was approved by the institutional review board at the University of Colorado at Boulder.

## RESULTS

As shown in **Table 1**, there were no significant differences in height, body weight, maximal oxygen consumption, or systolic and diastolic BP at rest between the groups. The majority of the participants had normal sinus rhythm while 4 participants had right bundle branch block at rest. ST segment depression was detected in 8 walkers (2 men, 6 women) and 4 runners (4 men). Walkers with ST-segment depression were older than other groups (*P*=.011). Maximal systolic BP during exercise was significantly higher in runners with ST-segment depression than those without (*P*=.024). This difference remained significant even after adjusting for sex differences using analysis of covariance (*P*=.045). There were no significant differences in maximal diastolic BP among groups.

**Table 1.** Physical characteristics of the participants and blood pressure responses during maximal exercise. Values are means±SD. ST, ST-segment depression;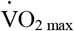, maximal oxygen uptake; SBP, systolic 0 blood pressure; DBP, diastolic blood pressure. ∗ p<0.05 vs. walker without ST; † p<0.05 vs. runner with ST.

|  | Walkers |  | Runners |  | P-value |  |  |
| --- | --- | --- | --- | --- | --- | --- | --- |
|  | Without ST | With ST | Without ST | With ST | Modality | ST | Modality x ST |
| M/F, n | 1/12 | 2/6 | 8/5 | 4/0 |  | NA |  |
| Age, yr | 68 ± 3 | 74 ± 4*† | 69 ± 5 | 67 ± 3 | .582 | .693 | .011 |
| Height, cm | 161 ± 10 | 158 ± 3 | 169 ± 9 | 176 ± 3 | .241 | .776 | .275 |
| Body mass, kg | 62 ± 11 | 53 ± 3 | 64 ± 13 | 76 ± 3 | .448 | .901 | .078 |
| $\dot{V}O_{2\text{ max}}$, ml/kg/min | 28 ± 4 | 31 ± 4 | 37 ± 6 | 38 ± 4 | .080 | .328 | .648 |
| SBP, mmHg | 120 ± 11 | 135 ± 12 | 122 ± 18 | 141 ± 17 | .298 | .065 | .760 |
| DBP, mmHg | 73 ± 8 | 72 ± 9 | 74 ± 12 | 74 ± 10 | .366 | .678 | .868 |
| *Maximal exercise* |  |  |  |  |  |  |  |
| Maximal SBP, mmHg | 188 ± 19 | 178 ± 21 | 187 ± 16 | 213 ± 26*† | .524 | .726 | .015 |
| Maximal DBP, mmHg | 79 ± 10 | 76 ± 11 | 76 ± 7 | 75 ± 7 | .310 | .291 | .739 |
| ΔSBP, mmHg | 68 ± 20 | 47 ± 23 | 62 ± 20 | 75 ± 7 | .834 | .978 | .022 |
| ΔDBP, mmHg | 7 ± 11 | 4 ± 8 | 1 ± 13 | 1 ± 13 | .407 | .214 | .701 |

## DISCUSSION

Exercise stress tests are typically performed on symptomatic patients and those at elevated risks of ischemic heart disease ^1,2^. In more recent years, these tests are increasingly recommended for masters athletes who regularly perform exercise training ^2^. In the sample of apparently healthy physically active older adults who were recruited for a research study to evaluate exercise economy ^7^, we found the coexistence of ST-segment depression and exaggerated BP response in older runners. These findings are consistent with the notion that the screening and assessment of cardiovascular risks should be extended to masters athletes who vigorously train regularly and are seemingly healthy.

The present study was not the first to demonstrate the coexistence of ST segment depression and peak systolic BP. A previous study reported that middle-aged long-distance runners with exercise-induced hypertension were found to have a higher prevalence of ST segment depression compared with those with normal exercise systolic BP ^2^. The present findings extend these research findings to older endurance-trained athletes. Taken together, these results suggest that the prevalence of ST-segment depression and exaggerated systolic BP response might be higher than previously thought in habitually trained adults.

In the present study, ST segment depression was not associated with the exaggerated systolic BP response in the group composed of walkers. In a low-risk group with a physically demanding occupation and without objective evidence of coronary artery disease, ST segment depression was not associated with systolic BP responses during exercise ^6^. Exaggerated systolic BP response appears to be much higher in prevalence in endurance-trained adults ^5^. The coexistence of ST-segment depression and greater peak systolic BP might be related to the exercise intensity that these athletes perform.

## ACKOWLEDGEMENTS

We thank Dr. Rodger Kram for contributing to early phases of this project.

## REFERENCES

1. Lauer M, Froelicher ES, Williams M, Kligfield P. Exercise testing in asymptomatic adults: a statement for professionals from the American Heart Association Council on Clinical Cardiology, Subcommittee on Exercise, Cardiac Rehabilitation, and Prevention. Circulation. Aug 2 2005;112(5):771–6. doi:10.1161/circulationaha.105.166543

2. Maron BJ, Araújo CG, Thompson PD, et al. Recommendations for preparticipation screening and the assessment of cardiovascular disease in masters athletes: an advisory for healthcare professionals from the working groups of the World Heart Federation, the International Federation of Sports Medicine, and the American Heart Association Committee on Exercise, Cardiac Rehabilitation, and Prevention. Circulation. Jan 16 2001;103(2):327–34. doi:10.1161/01.cir.103.2.327

3. O’Neal WT, Qureshi WT, Blaha MJ, Keteyian SJ, Brawner CA, Al-Mallah MH. Systolic Blood Pressure Response During Exercise Stress Testing: The Henry Ford ExercIse Testing (FIT) Project. J Am Heart Assoc. May 7 2015;4(5)doi:10.1161/JAHA.115.002050

4. Trenkwalder P, Dobrindt R, Plaschke M, Lydtin H. Usefulness of simultaneous ambulatory electrocardiographic and blood pressure monitoring in detecting myocardial ischemia in patients >70 years of age with systemic hypertension. The American Journal of Cardiology. 1993/10/15/1993;72(12):927–931. doi:10.1016/0002-9149(93)91109-U

5. Tanaka H, Bassett DR, Jr., Turner MJ. Exaggerated Blood Pressure Response to Maximal Exercise in Endurance-Trained Individuals*. American Journal of Hypertension. 1996;9(11):1099–1103. doi:10.1016/0895-7061(96)00238-5

6. Carlén A, Gustafsson M, Åström Aneq M, Nylander E. Exercise-induced ST depression in an asymptomatic population without coronary artery disease. Scandinavian Cardiovascular Journal. 2019/07/04 2019;53(4):206–212. doi:10.1080/14017431.2019.1626021

7. Ortega JD, Beck ON, Roby JM, Turney AL, Kram R. Running for exercise mitigates age-related deterioration of walking economy. PLoS One. 2014;9(11):e113471. doi:10.1371/journal.pone.0113471

